# Defining the microenvironment landscape of bladder cancer using highly multiplexed spatial genomic and proteomic analysis

**DOI:** 10.1101/675926

**Authors:** Jason W Reeves, Zhaojie Zhang, Zachary K Norgaard, Denise M Zhou, JingJing Gong, Yan Liang, Subhasree Das, Sarah E Warren, Manav Korpal, Margaret L Hoang, Joseph M Beechem, Pavan Kumar, Victoria Rimkunas

## Abstract

Muscle-invasive bladder cancer (MIBC) is an aggressive disease with limited therapeutic options. PD-1 pathway targeting immunotherapies have been approved to treat advanced bladder cancer, but most patients exhibit primary resistance, suggesting that immune evasion mechanisms exist. The PPARγ pathway has been identified as a potential therapeutic target in MIBC that is associated with reduced CD8+ T-cell infiltration and increased resistance to immunotherapies. We comprehensively profiled the tumor microenvironment (TME) in formalin-fixed, paraffin-embedded (FFPE) tissues from a cohort of PPARγ^high^ (n=13) and PPRARγ^low^ (n=12) MIBC, integrating bulk gene expression, targeted mutation sequencing, immunohistochemistry and multiplex spatial profiling of RNA and protein expression on the GeoMx™ Digital Spatial Profiling (DSP) platform. Molecular subtyping was consistent between traditional methods and GeoMx profiling, and, in this cohort, we observed little evidence of spatial heterogeneity in tumor subtyping. The previously characterized T-cell exclusion phenotype of PPARγ^high^ MIBC was recapitulated on the GeoMx platform and was further extended to show that this is a general phenomenon across immune cell types, supporting potential combination of PPARγ inhibition with ICIs. Furthermore, we found that while immune cells were excluded from PPARγ^high^ tumors, the stromal compartment from these tumors was not significantly different than those PPARγ^low^ tumors. By preserving spatial relationships during the GeoMx analysis, we also identify a novel association between lower immune cell expression in the tumors and higher expression of β-catenin in the stroma, and differential expression of other WNT pathway members associated with PPARγ activity.

**One Sentence Summary:** A new method for capturing tumor-immune signaling in FFPE tissues explores how the PPARG signaling axis is associated with immune cell exclusion in bladder cancer.

## Introduction

Muscle invasive bladder cancers (MIBCs) are biologically heterogenous and have widely variable clinical outcomes and responses to conventional chemotherapy(*1–4*) and immune checkpoint inhibitors(*5*). Although responses can be durable, only 20-30% of patients with MIBC will respond to immune checkpoint inhibitors (ICIs), arguing for the identification of novel drug targets and need for a deeper understanding of predictive biomarkers to enrich for responses to current standard of care therapies(*6, 7*). Multiple studies have identified distinct RNA expression subtypes within MIBC termed “luminal” and “basal-like”(*8*), each of which has unique gene expression patterns which have important implications for disease management with conventional chemotherapy(*9–11*) and immunotherapies(*12*). Response to immune checkpoint inhibitors in urothelial bladder cancer has been associated with several potential biomarkers including tumor mutation burden, tumor molecular subtype and PD-L1 expression on CD8+ tumor infiltrating lymphocytes (TILs) and other immune cells(*7, 13–15*).

Patients with the luminal I subtype rarely respond to ICIs(*14, 15*), suggesting the existence of one or more immune escape mechanisms. We previously reported that the PPARγ/RxRa pathway constitutes a tumor-intrinsic mechanism underlying immune evasion in MIBC (*6, 16*). In tumors with high PPARγ pathway activity, CD8+ T-cell infiltration appears to be impaired through chemokine suppression, though additional mechanisms may also impact T-cell infiltration. These findings suggest that pharmacological inhibition of PPARγ may induce sensitivity to immunotherapies. To further our understanding of tumor-immune escape mechanisms in MIBC, analysis of both the tumor and the surrounding microenvironment is necessary. Comprehensive interrogation of the interaction between these two compartments has proved challenging thus far because most of the foundational work in MIBC has relied on commonly available assays such as single-plex immunohistochemistry (IHC) assays or bulk RNA expression and mutation profiling. With limited availability of tumor tissues during clinical trials, highly multiplexed and quantitative platforms compatible with small formalin-fixed paraffin-embedded (FFPE) tissues are necessary to deliver a comprehensive understanding of the tumor-immune landscape.

Historically, protein assessments of the TME have been limited to PDL1 and CD8 single plex IHC staining, which is primarily driven by the observation that in urothelial carcinoma and other cancers, outcomes related to ICI treatment appear particularly favorable in patients with high PD-L1 expression(*7, 13, 14, 17, 18*). But these assays for PD-L1 have been developed for individual therapies, using varying antibody clones, staining protocols, scoring algorithms and cut-offs, which makes standardization across multiple studies difficult (*19, 20*). In addition, conventional IHC has significant limitations due to relatively low sensitivity for poorly antigenic or expressed targets, difficulty in colocalizing stains, and the subjective and semi-quantitative nature of pathologists’ interpretations. While significant progress has been made in multiplex IHC technologies, significant challenges remain for the field. Immunofluorescence (IF) platforms such as Ultiview, Multiomyx and Opal can provide quantitative analysis of multiple stains from a single slide, but the analysis is quite complex, time consuming and costly, and except for Multiomyx, most multiplex IHC technologies are limited to 4-7 analytes per slide, which is insufficient for fully characterizing the TME.

Given the limitations of IHC, gene expression profiling is an attractive alternative allowing simultaneous profiling of hundreds to thousands of transcripts from as little as 50 ng of RNA. In high quality samples, this can be extracted from a single FFPE slide, although the RNA yield varies drastically by tissue sample size, tumor type, and age of the slide. RNAseq and NanoString® nCounter™ direct hybridization are two bulk gene expression methods that have routinely been used for studying the TME, resulting in identification and validation of gene signatures like the tumor inflammation signature (TIS)(*21, 22*) showing correlation with response to ICIs in other cancers(*19, 23*). The breadth and scale of gene expression analysis has been foundational in oncology, but these methods lack the information conferred by spatial resolution of the signal. Previous studies in colorectal cancer and melanoma have highlighted the critical role that the spatial localization of the immune compartment plays in patient outcome(*23–25*), while the role of the spatial and temporal distribution of CD8 TILs and PD1 expression has been suggested to be predictive of response to ICIs (*17, 18*). Spatial resolution of RNA biomarkers can be retained with technologies such as RNAscope (in situ hybridization, ACD) but similar to multiplex IHC, only 1-3 analytes per FFPE section can be measured. A higher resolution and multiplexing can be afforded by a number of methods in fresh frozen tissues (*26–28*), but these require specialized workflows not frequently compatible with clinical trials nor are such approaches applicable to archived FFPE tissues.

In this report, we describe the application of multiple bulk analysis and spatial multiplexed technologies to characterize the TME in a cohort of MIBC FFPE tumor samples, using tissue stored in FFPE blocks previously used to characterize the CD8 immune-exclusion phenotype associated with PPARγ activity (*6*). For bulk profiling, we applied both RNA sequencing and NanoString PanCancer IO 360 gene expression readouts, as well as a targeted sequencing panel for common cancer-associated mutations. To resolve spatial interactions, we processed samples using the NanoString GeoMx platform using both multiplexed RNA in situ hybridization (ISH) and a multiplexed antibody cocktail, while using traditional single-plex IHC or ISH for validation purposes. This platform uses UV photocleavable oligonucleotide tags conjugated antibodies or RNA-ISH probes to interrogate targets expression in a tissue section. The oligonucleotides are released from specific regions of interest (ROIs) via exposure to UV light, in paths that are controlled by a digital micromirror array (*29, 30*). The goals of this study were to interrogate the molecular composition of the TME of MIBC across complementary technologies, and to characterize the ability of different platforms to profile clinical samples where tissue is limited. We also explore the role of tumor-stromal interactions within this cohort to expand our understanding of potential ICI resistance mechanisms and opportunities for therapeutic combinations in advanced bladder cancer.

## Results

### Bulk and spatial approaches for understanding tumor-stromal interaction in MIBC

To investigate the interplay between tumor molecular subtype, PPARγ signaling, and the composition of the TME, we profiled a previously collected cohort of 25 samples from advanced stage MIBC for which PPARγ pathway activity and CD8 staining had been previously characterized (Table 1, Supplementary Table 1) (*6*). These samples had a higher proportion of male patients, a balance of samples with or without lymph node involvement, and with most samples being late stage, though 5 of the 13 PPARγ^high^ tumors were stage I-II. PPARγ^high^ samples had higher tumor cellularity on average whether estimated on either the original H&E (p = 0.04, Welch’s t-test, Supplementary Table 1) or IF channels from RNA (p < 0.008, Welch’s t-test, Supplementary Table 1). The workflow used to characterize this cohort included a multifaceted approach of combining traditional bulk sequencing and expression assays with spatial profiling by IHC/ISH assays and multiplexed protein and RNA quantification on the NanoString GeoMx platform (Figure 1A).

**Table 1.** Clinical, genomic, and TME characteristics of MIBC cohort. *median age and range shown. ^†^Estimated from RNA-ISH IF Pan-CK staining

| | | PPAR$\gamma^{\text{high}}$<br>(n = 13) | PPAR$\gamma^{\text{low}}$<br>(n = 12) |
| --- | --- | --- | --- |
| **Stage** | I | 2 | 0 |
|  | II | 3 | 0 |
|  | III | 5 | 5 |
|  | IV | 3 | 7 |
| **Age\*** |  | 58 (44-75) | 67 (43-72) |
| **LN Status** | Negative | 10 | 6 |
|  | Positive | 3 | 6 |
| **Gender** | Female | 1 | 3 |
|  | Male | 12 | 9 |
| **Genomic Subtype** | Basal | 0 | 8 |
|  | Luminal | 13 | 4 |
| **CD8 TILS** | < 1% | 9 | 3 |
|  | > 1% & < 5% | 3 | 4 |
|  | > 5% | 1 | 5 |
| **Tumor Cellularity<sup>†</sup>** | < 60% | 4 | 7 |
|  | > 60% | 9 | 2 |
| **GeoMx Profiling** | Antibody | 13 | 10 |
|  | RNA-ISH | 10 | 9 |

**Fig. 1.**
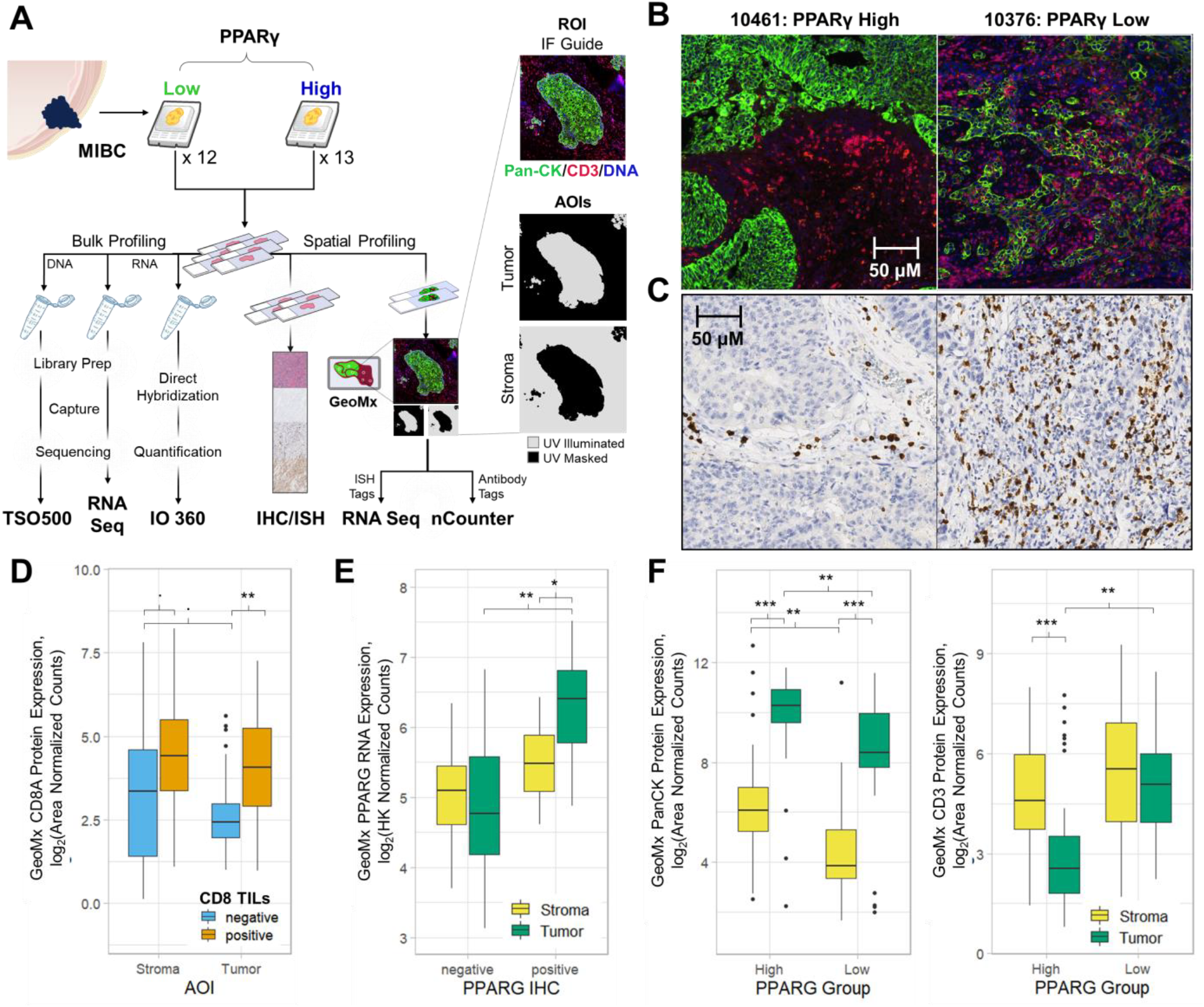
Study overview and qualification of GeoMx readouts **(A)** Patient samples analyzed using separate workflows for bulk and spatial profiling. Samples were prepared for multiple workstreams for bulk of DNA and RNA and spatial profiling of RNA and protein expression. GeoMx analysis involves selection of regions of interest (ROIs) based on whole-slide IF and subsequent UV illumination of specific paths into areas of illumination (AOI, gray) based on Pan-CK staining by IF to collect oligo tags for downstream quantification. **(B)** Example IF guide images placed for selected ROIs during GeoMx workflow and **(C)** related CD8 IHC images of the same ROI for either a representative PPARγ^high^ patient (left) or PPARγ^low^ patient (right) **(D-F)** Example validation of concordance between GeoMx profiling and **(D)** IHC of CD8 TILs, **(E)** PPARG expression by ISH, or **(F)** between guide markers and related antibody probes (Pan-CK, left; CD3, right). Nominal significance shown for mixed effect model, denoted as. for p < 0.1, * p < 0.05, ** p < 0.01, *** p < 0.001.

For each sample, the GeoMx workflow involved selection of regions of interest (ROIs), which can be quantitated using downstream sequencing or nCounter readouts. In this study, we selected 5-6 ROIs per patient by focusing on areas of the slide with both tumor-rich and stomal areas within the same ROI using Pan-CK and CD3 IF staining to assess tumor and stromal content, avoiding regions of necrosis or tissue detachment (Supplementary Figure 1). Examples of multiplexed IF images used to guide UV illumination are shown based on PPARγ status in (Figure 1B). Custom UV illumination masks were designed to create areas of illumination (AOIs) that result in photocleavable tags being released specifically in either tumor (Pan-CK+) or stromal (Pan-CK-) compartments, a process called auto-segmentation (*29, 30*) (Figure 1A). 23 samples of the cohort were profiled using multiplexed antibodies, collecting tags from 273 AOIs. Two samples failed due to tissue detachment during processing. 19 samples were profiled with multiplexed RNA-ISH collecting tags from 222 AOIs, after 3 samples failed due to tissue detachment. The tag oligonucleotides were quantified using either the nCounter platform, for antibody tags, or with next generation sequencing (NGS), for RNA-ISH tags. GeoMx measured expression of 40 proteins or 157 RNA transcripts simultaneously, using a single five-micron FFPE slide for each type of analyte. Of the AOIs profiled by RNA-ISH, 13 were flagged as having high background and removed from analysis (Supplementary Table 2, Supplementary Figure 2). 29 out of 38 protein targets stained were detected above the expression of the two IgG control antibodies (Supplemental Figure 1, Supplementary Data File 1). Similarly, for RNA-ISH readout the negative control probes were on average expressed at the 12^th^ percentile (±6%, σ_x_) of all probes tested after excluding samples with high background (Supplementary Figure 2, Supplementary Data File 2).

In addition to GeoMx analysis, sections were stained for CD8 and granzyme B by single-plex IHC. IHC images of CD8 expression from the same block, though not serial section, are shown for comparison with the IF images used to guide ROI selection (Figure 1C). CD8A was confirmed to be higher in tumor AOIs that stained positive for the CD8 TILs presence (p = 0.006, mixed effect model) and trended towards higher expression in stromal AOIs (p = 0.057, mixed effect model, Figure 1D). Since CD3 was not used to create UV illumination masks but rather to help guide ROI selection, CD8 expression within tumors AOIs was attributed to CD3+ cells present in tumor areas observed by IF in samples. Similarly, PPARG expression level detected by RNA-ISH GeoMx was highest in tumor AOIs from PPARG+ tumors identified by IHC (p < 0.05, mixed effect model, Figure 1E). Expression of markers associated with IF staining guide followed expected patterns based on IF images, with Pan-CK expression being significantly higher in tumor AOIs than stromal AOIs (p < 0.001, mixed effect model, Figure 1F). Finally, CD3 expression was more complex, with detection at equivalent levels in stromal AOIs, but significantly lower expression in tumor AOIs specifically from PPARγ^high^ tumors. This is consistent with the previous described exclusion phenotype(*6*), as CD3 IF staining was not used during auto-segmentation.

### Tumor molecular subtyping and intrinsic signaling of PPARγ on GeoMx

Since molecular subtype has been implicated in response to multiple therapies, it was important to understand the distribution of samples in this cohort and whether GeoMx profiling could classify the MIBC cohort into luminal or basal tumor intrinsic subtypes. To provide a baseline for analysis, we classified samples based on bulk RNA sequencing data into basal or luminal genomic subtypes(10). 17 samples were classified as luminal (68%) and 8 as basal (32%, Table 1), with membership skewed based on PPARγ status as previously reported from analysis of TCGA(*6*). CD44, a basal MIBC marker, was the only subtype marker included in both protein and RNA-ISH assays. In the protein assays, we calculated the receiver-operating characteristic (ROC) curve and found that CD44 was more predictive in tumor AOIs (AUC = 0.89) than stromal AOIs (AUC = 0.66, Figure 2A).

**Fig. 2.**
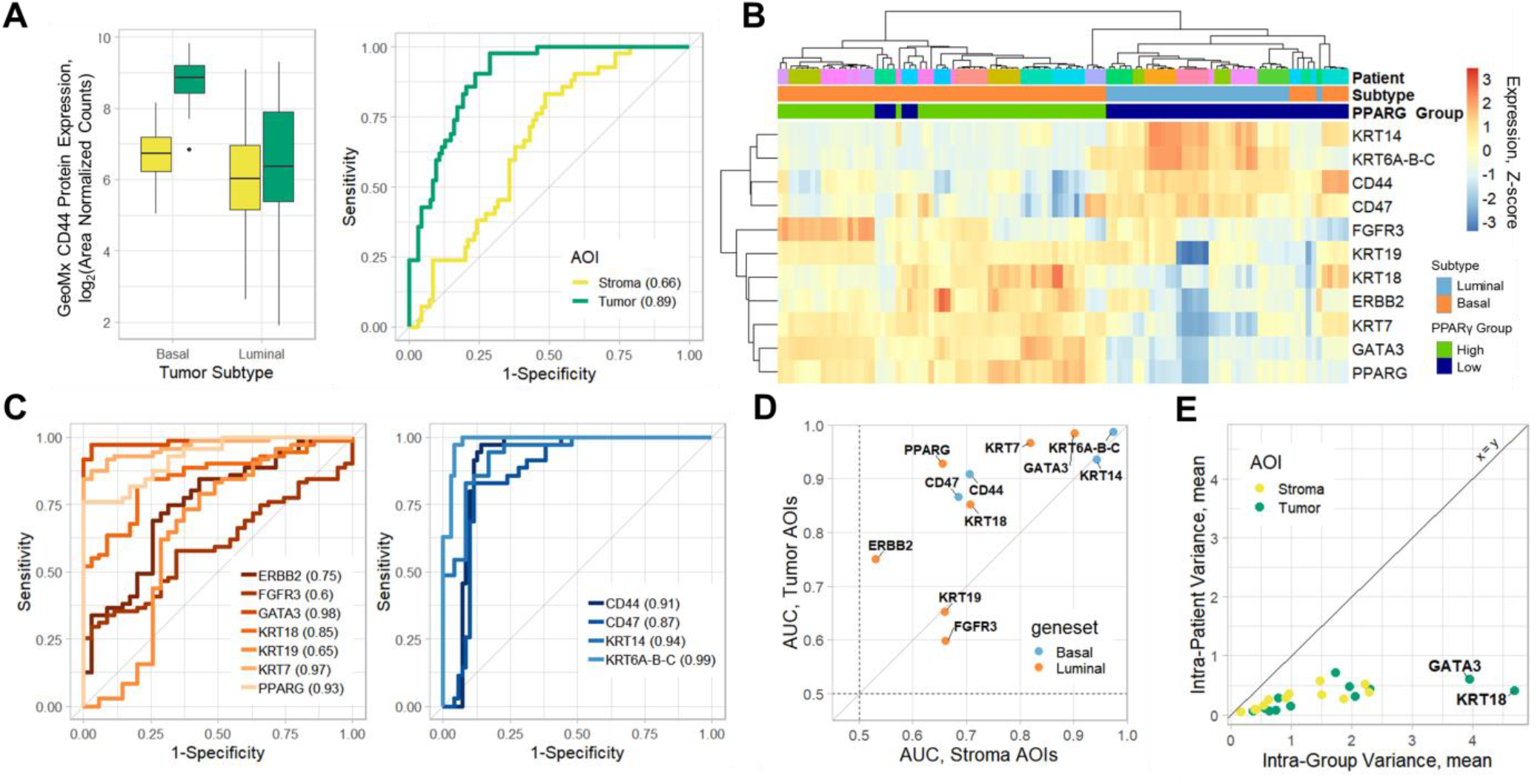
Tumor intrinsic subtyping (**A**) Analysis of protein expression of CD44 as a marker for tumor subtype, per AOI. ROC curves shown for tumor and stromal AOIs analyzed separately. AUC values shown for each AOI type. **(B)** Unsupervised hierarchical clustering of subtyping probes included in RNA-ISH GeoMx analysis. Annotations show patient ID, RNA sequencing identified subtype (orange = luminal, light blue = basal), and PPARG expression group (dark blue = low, green = high) **(C)** ROC analysis of MIBC subtype classification for RNA-ISH markers measured in tumor AOIs. **(D)** Comparison of AUCs from ROC analysis of tumor or stromal AOIs from C. **(E)** Comparison of average variance within subtype or patient samples

For RNA-ISH analysis, we first qualitatively explored the performance of probes to identify each subtype by unsupervised hierarchical clustering across the tumor AOIs. Subtype markers cluster tumor AOIs into basal and luminal tumors as classified by bulk RNA sequencing with AOIs within sample as well (Figure 2B). While the clustering was not completely resolved by subtype, the targeted panel for ISH only contained 12 subtyping markers previously described, and it is possible that profiling additional markers would provide better resolution.

To quantify the predictive power of RNA-ISH profiling to classify samples, we calculated a ROC curve for each gene. We found that markers for subtype generally performed well individually in tumor AOIs (Figure 2C), though basal markers were more consistently predictive than luminal markers. Consistent with the protein analysis, these markers were less predictive in stromal AOIs, although some keratins remained highly predictive (Figure 2D). Testing of intratumor heterogeneity among subtyping markers revealed that the average variation in expression was always higher within a given subgroup than within a sample, and that variance was equivalent between tumor and stroma AOI (Figure 2E). This suggests that intra-tumoral heterogeneity in tumor-intrinsic signaling is not likely a major factor in the AOIs sampled in our cohort.

### Bulk analysis masks expression in stromal compartments

Based on the observation that subtyping genes were more predictive in the tumor AOIs, we explored the source of expression derived from bulk assays. Tumor purity, and its impact on expression profiling, has been studied in the context of the TCGA, and bladder cancers were found to have higher signal from the stroma than other many cancers (*31*). However, it’s unclear if these bulk studies reflect subtype-specific TME structural differences, or true stromal expression patterns. Therefore, we compared our RNA-ISH results to the bulk RNA profiling by the nCounter PanCancer IO 360 assay as these were the most similar in terms of probe design and analysis workflow. We tested whether either type of AOI, Pan-CK+ tumor or Pan-CK-stroma, was better correlated with bulk profiling and tested the impact of tumor cellularity on the correlation observed with either AOI type. We were able to profile 108 genes across both platforms. We found that tumor AOIs had significantly higher correlation with bulk expression within a given patient across all shared genes in the two platforms (p = 4.8e-4, Welch’s t-test, Figure 3A). Patients with higher tumor cellularity as estimated from the whole-slide IF showed a slight, though insignificant, increase in correlation between bulk tumor expression and tumor AOIs (p = 0.11, Welch’s t-test), and no appreciable difference within stromal AOIs (p = 0.26, Welch’s t-test).

**Fig 3.**
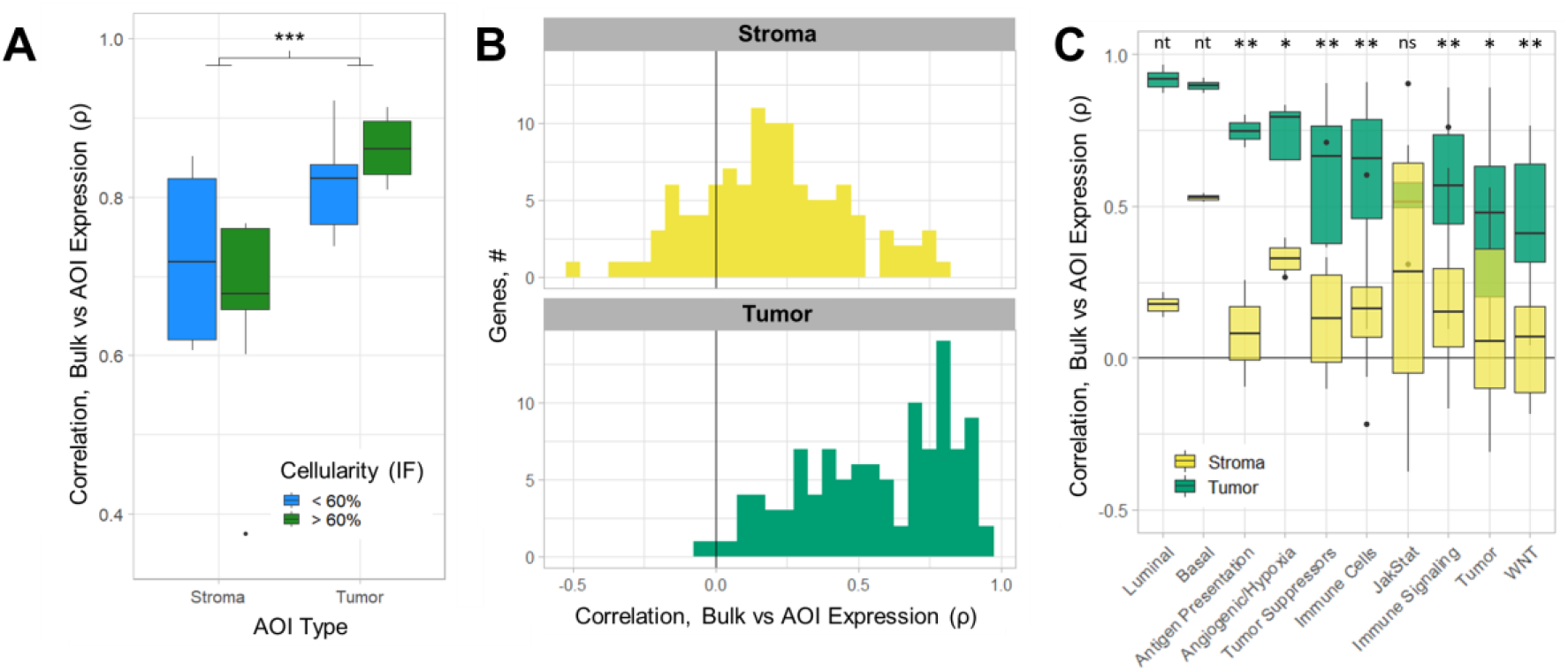
Bulk profiling masks expression patterns observed in the stroma **(A)** Correlation of gene expression observed in IO360 profiling and RNA-ISH GeoMx profiling. Correlations were calculated between bulk samples and each AOI from the same sample and then separated by AOI type and tumor cellularity estimated from the IF image used to select ROIs. Significantly higher (p < 0.001, Welch’s t-test) correlation observed in tumor AOIs than Stromal AOIs. **(B-C)** Gene-based calculations of correlation based on average AOI expression for a patient within each AOI type, showing correlations across all genes based on **(B)** AOI location (Stroma, top; tumor, bottom) and **(C)** associated gene set information. Significance (p, Welch’s t-test) indicated by * < 0.05, ** < 0.01, ns > 0.05, nt for not tested when *n*_*genes*_ < 3

Similarly, we found that, on a gene level, expression was more highly correlated with bulk expression in tumor AOIs (mean ρ = 0.56, Spearman) than stromal AOIs (mean ρ = 0.20; p < 1e-16, Welch’s t-test, Figure 3B, Supplementary Figure 3). Higher average gene expression in tumor AOIs was also moderately associated with higher correlation with bulk expression(ρ = 0.55, Supplemental Figure 3). When assessing genes from different pathways or gene sets, genes that were part of tumor-intrinsic subtyping gene sets were the most highly correlated between tumor AOIs and bulk expression (Figure 3C, Supplementary Figure 3), though most gene sets followed the same pattern. Collectively, these data suggest that bulk profiling of bladder cancer more closely reflects the tumor compartment and under-represents expression from the surrounding stroma.

### Spatial organization of TILs in MIBC associates with PPARγ expression

To further explore the biologic basis of immune exclusion in our cohort, we performed Gene Set Enrichment Analysis (GSEA) on the RNA-sequencing data with cancer hallmark signatures (*32*). Immune related hallmarks, such as inflammatory response, IL6/JAK/STAT3 signaling, IL3/STAT5 signaling and TNFA signaling are enriched in the PPARγ^low^ group (Figure 4A), while PPARγ signaling-related signature such as Oxidative Phosphorylation and Fatty Acid Metabolism were anticorrelated with this group (Supplementary Data File 3). In addition, results from IO 360 profiling of the TIS (17,20) found that TIS was anticorrelated with PPARG expression (ρ = −0.66, Spearman, Figure 4B). TIS expression was positively correlated with CD8+ TIL infiltration (ρ = 0.72, Spearman) and PD-L1 expression (ρ = 0.91, Spearman), while PPARG RNA expression was inversely associated with both (ρ = −0.43, −0.73 respectively, Spearman). In addition to TIS, nearly all immune-related signatures calculated from IO 360 were lower in PPARγ^high^ tumors (Supplementary Data File 3), suggesting this exclusion extends beyond the T-cell infiltration previously characterized (Figure 4C).

**Fig 4.**
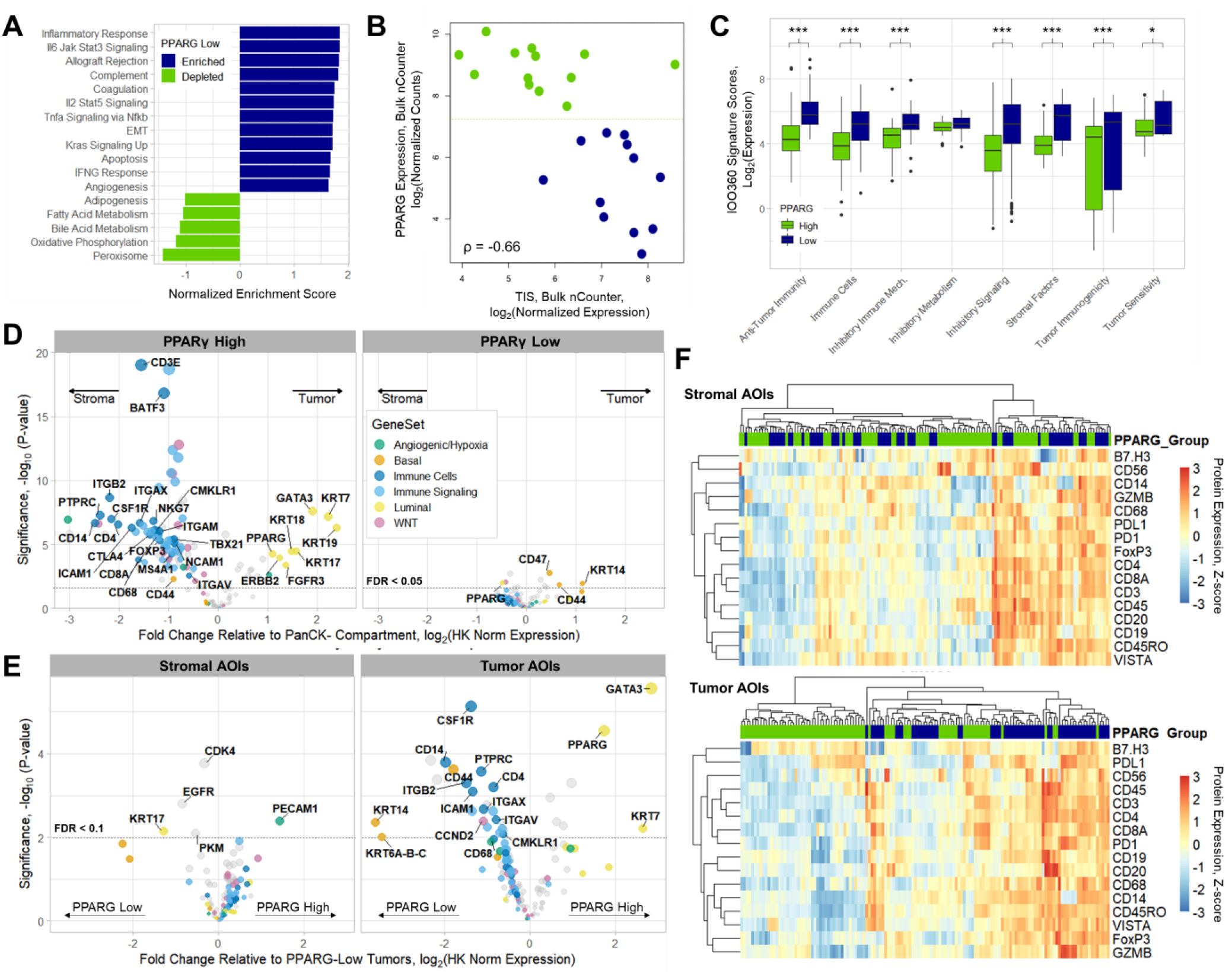
Phenotypic analysis of the tumor microenvironment in MIBC **(A)** Gene set enrichment analysis results from bulk RNA-sequencing analysis indicative of high immune-cell related signaling detection in PPARγ^low^ tumors **(B)** PPARγ expression level vs tumor inflammation signature (TIS), colored base to PPARγ status from IO360 analysis (high = green, low = dark blue) **(C)** IO360 Signatures score expression based on PPARγ status, binned by type of signature. Significance (p, Welch’s t-test) indicated by * < 0.05, ** < 0.01, *** < 1-e3 **(D)** Differential expression analysis GeoMx RNA-ISH data shows stromal enrichment of immune cell and immune signaling genes in PPARγ^high^ tumors (left), while no significant difference is observed in PPARγ^low^ tumors (right). Genes colored based on gene sets, labelled for immune cells or subtyping markers for FDR < 0.05 **(D)** Differential expression analysis GeoMx RNA-ISH data based on PPARG status in each type of AOI. Genes colored based on gene sets, labelled for immune cells or subtyping markers for FDR < 0.1 in Tumor AOIs, all genes with FDR < 0.1 labeled in stromal AOIs (**F)** USHC of immune cell markers from antibody analysis of either stromal AOIs (top) or tumor AOIs (bottom).

To identify immune exclusion and differentiate this from the tissue being devoid of immune cells (i.e. an immune desert), we compared the expression of protein or RNA markers between tumor and stromal AOIs, while controlling for inter-patient variability in expression in either PPARγ^high^ or PPARγ^low^ tumors. We found that PPARγ^high^ tumors, but not PPARγ^low^ tumors, exhibited significant differential expression of immune cell markers. PPARγ^high^ samples had higher expression in the stromal AOIs after accounting for multiple testing (FDR < 0.05, Figure 4D, Supplementary Table 3). Markers enriched in the stromal compartment included numerous cell type markers, including lineage specific markers such as CD8A, CD56/NCAM1, ICAM1, CD14, CD68, and CSF1R. Furthermore, many immune signaling markers including cytokines and chemokines measured as part of the RNA-ISH DSP were found to be restricted to the stroma in PPARγ^high^ tumors, as anticipated since immune trafficking is impaired in these tumors. In contrast, while nominal differential expression was observed between tumor and stroma AOIs for PPARγ^low^ tumors, no immune cell markers reached significance after adjusting for multiple testing (FDR < 0.05).

While the immune exclusion phenotype has been previously demonstrated, it was unclear whether the composition of immune cells in the stroma was different between the two subtypes of the disease. We examined this by comparing the expression of markers in the stromal compartments between PPARγ^high^ and PPARγ^low^ tumors. Under this analysis, few markers reached significance after adjusting for multiple test correction in either the protein or RNA dataset, even while using a less stringent false discovery rate (Figure 4E, FDR < 0.1). We also ran unsupervised hierarchical clustering of antibodies related to immune cell types or signaling from stromal AOIs and compared the results to similar analysis of tumor AOIs (Figure 4F). Under this analysis we found that PPARγ groups cluster based on tumor AOIs expression, while stromal AOIs did not have a similar clustering structure. Moreover, under traditional IHC analysis we did not see significant differences in GZMB expression based on PPARγ expression (Supplemental Figure 4), suggesting no differential impact on the cytolytic potential in cells neighboring the tumor. Taken together these results suggest that PPARγ expression does not drive changes in composition of the immune cells outside the tumor, though its expression is related to their exclusion.

### High tumor mutation burden found in PPARγ^high^ patient sample with high TIL presence

Despite the immune-exclusion mechanisms associated with PPARγ expression, we identified one patient sample, 10182, expressing high PPARγ while also having high TIS and increased TIL presence by IF or IHC staining (Figure 5A-B). This was reproduced in GeoMx analysis, showing that CD3 protein expression was similar between tumor and stromal AOIs, and observed at levels similar to those of PPARγ^low^ tumors (Supplemental Figure 5). One possible explanation for an inflamed TME is a high tumor mutation burden which others have shown may be an orthogonal metric for immune activation from general tumor infiltration and inflammatory signaling (*23*). As this would not be directly captured in any of the other assays run, we profiled samples on the TSO-500 targeted sequencing panel, comprising 500 commonly altered genes in cancer. From this assay, tumor mutational burden (TMB) was estimated based on the rate of nonsynonymous mutations observed in the panel targets.

**Fig 5.**
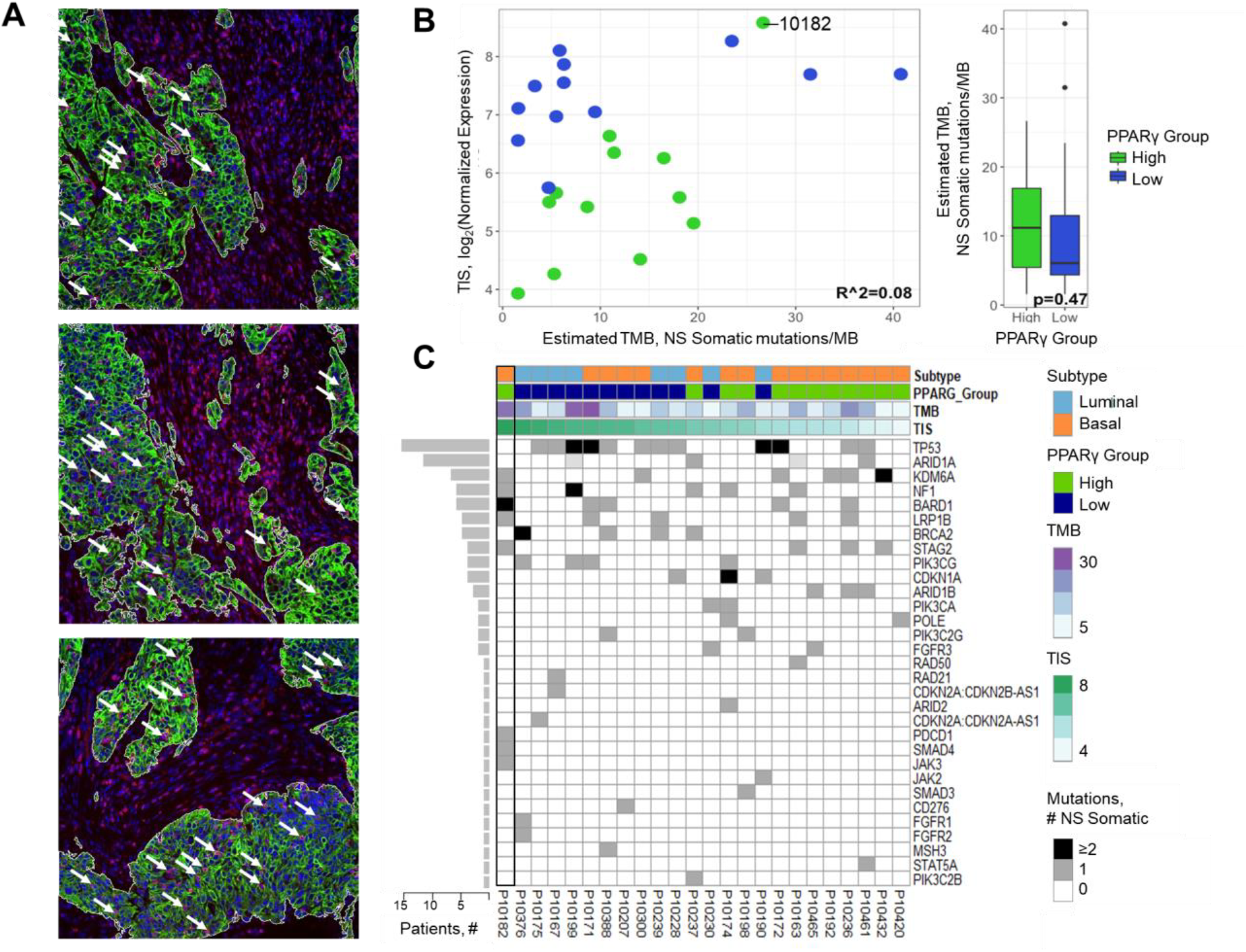
Tumor mutations associated with high TILs in patient 10182 **(A)** IF images of ROIs collected from sample 10182. Markers stained include Pan-CK (green), CD3 (red), and nuclear stain (blue). Arrows indicate CD3+ TILs within Pan-CK AOIs. **(B)** TMB analysis of bulk samples vs TIS or PPARG group, highlighting sample 10182 **(C)** Oncoprint of mutations across samples showing tumor-intrinsic subtype, PPARγ expression, TMB counts, and TIS.

While we found that TMB was not significantly different based on PPARγ expression, we found that sample 10182 had the highest TMB within PPARγ^high^ tumors (Figure 5B-C). We also found that in addition to a high mutational load, there were also specific mutations in genes related to immune signaling pathways including PDCD1, STAT3, and SMAD4 (Figure 5D). Of particular note, the SMAD4 mutation, T338I, was found within the MH2 functional domain and was predicted to be oncogenic (0.92, Cscape score) (*33*). Signaling within the TGFβ pathway, which is may be impacted by this mutation, has been reported to impact TIL presence and response to ICIs (*34*). Together, both TMB and specific mutations may represent compensatory mechanisms allowing TIL invasion despite PPARγ signaling in this patient.

### Stromal β-catenin signaling inversely associated with TILs identified by spatial genomics approaches

As TIL exclusion was related to tumor-intrinsic PPARγ, we wanted to further explore interactions between the tumor and stromal compartment that would be missed by bulk profiling or subgroup specific analysis. Because we profiled samples with matched tumor-stroma pairs of AOIs (n = 95, paired AOIs after QC), we designed a framework to test whether the tumor-stroma relationship between any sets of genes on the RNA-ISH panel was not observed in their native compartment and might be the result of paracrine or juxtacrine signaling (Figure 6A). Specifically, we calculated whether the tumor expression of a gene, *T*, was correlated with stromal expression another gene, *S* (|ρ(*T,S*)| > 0.5, Spearman), but not when comparing expression in the same compartment (|ρ(*T,T*; *S,S*)| < 0.2, Spearman). This allows us to detect potential interactions that subgroup-based analyses might miss. For example, we found that CTNNB1 (β-catenin) expression in stromal AOIs was anticorrelated with PTPRC (CD45) expression in the tumor AOIs (ρ = −0.56), but not in any other combination of AOI comparisons (Figure 6B).

**Fig 6.**
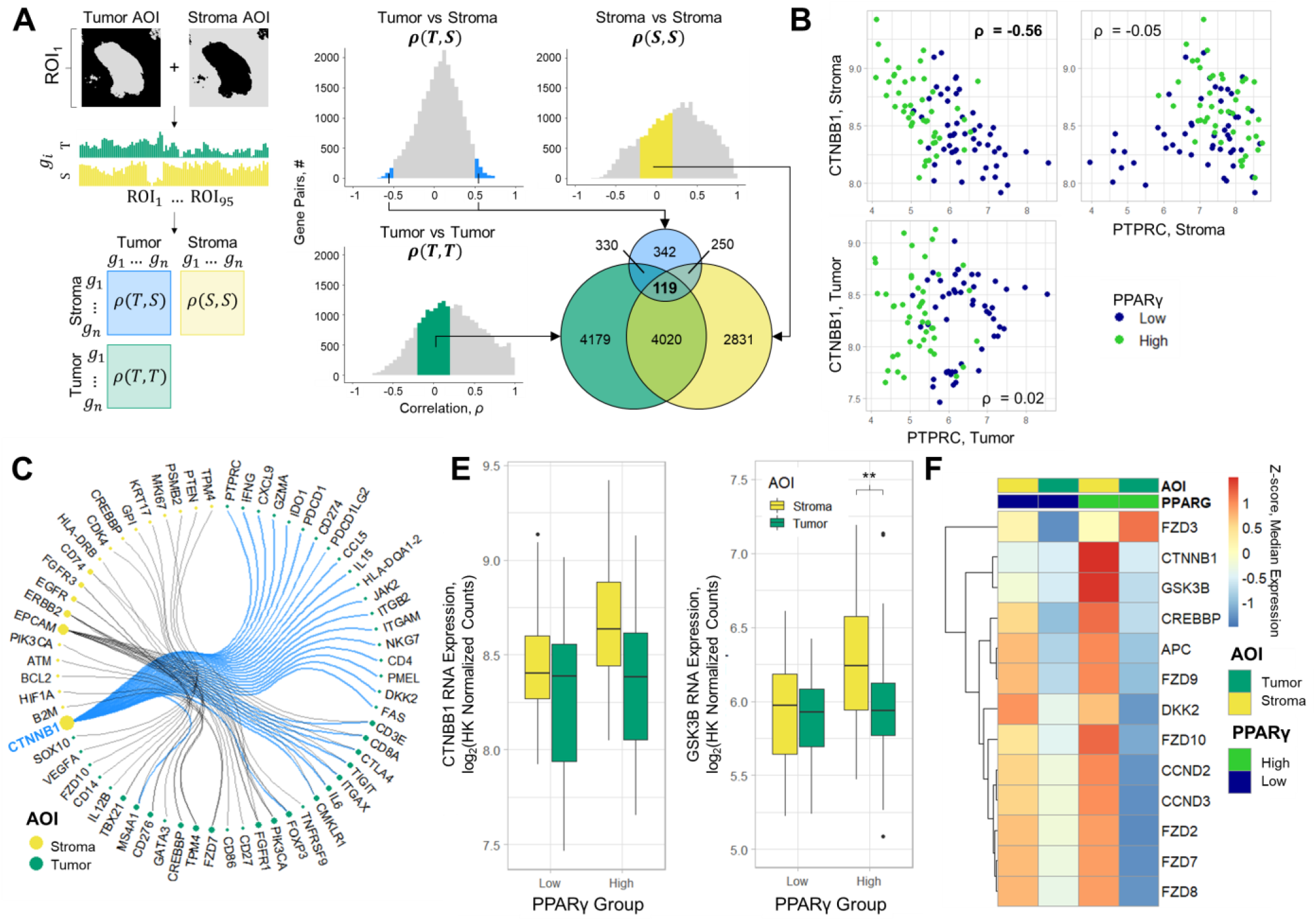
Spatial analysis identifies a stromal WNT-β-Catenin axis associated with immune cells **(A)** RNA-ISH tag expression from AOIs (gray areas, left) was used to calculate the correlation coefficient (ρ, Spearman) for expression values for all pairs of genes across different combinations of types of AOIs measured across 95 ROIs. Potential paracrine associations (n = 119) were identified by selecting genes with high correlation between expression in tumor and stromal compartments (|ρ(*T,S*)| > 0.5), while also having minimal correlation within the same compartment type (|ρ(*T,T; S,S*)| < 0.2). **(C)** Expression of CTNNB1 (β-Catenin) and PTPRC (CD45) based on compartment of measurement, demonstrating the specific correlation between stromal β-catenin and tumor expression of PTPRC **(D)** Map of identified tumor-stroma interactions. Lines indicate tumor-stroma specific correlations, with pairs that include stromal CTNBB1 highlighted in blue. Nodes colored by AOI of expression, size based on number of interactions identified. **(E)** Expression of β-catenin and GSK3B based on PPARγ expression and AOI. ** FDR < 0.01 **(F)** Heatmap of WNT/β-catenin pathway members in each compartment based on PPARγ expression.

Across all genes on the panel, we found that 119 pairs of genes exhibited this type of tumor-stroma specific correlation. As these interactions could be driven by patient-specific expression, we followed this analysis with an additional regression accounting for the patient from which an ROI was collected. Of the identified gene interactions, 65 were significant after accounting for patient-specific expression and multiple testing (FDR < 0.05, linear mixed effect model, Supplementary Table 3). The most frequently identified gene was CTNNB1 expression in stromal AOIs. CTNNB1 was anticorrelated with tumor expression of 28 other genes, most of which were noted to be immune cell markers or signaling molecules (Figure 6C). The next most common interaction partner identified was stromal expression of EPCAM, which had 12 significant interactions. While the antibody panel contained β-catenin and several of the immune cell targets identified in this analysis, we were unable to validate this finding in the protein GeoMx data (Supplementary Table 3). We attribute this to the fact that β-catenin localization, reflective of β-catenin activity, is not captured during GeoMx analysis, confounding our interpretation of the antibody data. Furthermore, analysis of bulk expression showed that tumor expression of CTNNB1 was correlated with bulk profiling (ρ = 0.69), suggesting accurate measurement of the target since CTNNB1 was observed to have higher expression in the stroma than in tumor compartment, though this was not significant after adjusting for multiple testing and patient-specific expression (FDR = 0.15, mixed effect model, Figure 6D).

To better understand the potential role of the β-catenin signaling axis in these tumors, we returned to the expression of other genes on the RNA-ISH panel which were not specifically identified by the previous analysis. We noted that several genes in the WNT/β-Catenin signaling pathway were identified as significantly enriched in the stroma of PPARγ^high^ tumors (Figures 3D). One of the members of this pathway, GSK3B, had similar patterns of expression across sample types as β-Catenin itself, and was specifically higher in PPARγ^high^ stroma (FDR = 6.4e-3, mixed effect model, Figure 6D). Of the 6 frizzled-family receptors on the RNA-ISH panel, all except FZD3 had higher median expression within the stroma of PPARγ^high^ tumors (Figure 6F). Additionally, other WNT/β-Catenin targets, such as CCND2/3, CREBBP, and APC were all similarly highest in stroma of PPARγ^high^ tumors, though expression was observed in PPARγ^low^ stroma as well. The differential expression of these genes appears to be driven by very low expression within the tumor compartment.

## Discussion

The study we report here follows the analysis of a cohort of advanced stage MIBC. In this setting, understanding the molecular underpinnings of a sample is critical to better predicting how they will respond to therapy(*10*), especially ICIs(*12, 13*). To better characterize the spatial arrangement of key molecular characteristics such as tumor-intrinsic subtype, TIL exclusion and composition, and tumor-stroma interaction we leveraged multiple traditional bulk and spatial platforms in conjunction highly multiplexed profiling by GeoMx within FFPE tissues. This allowed us to deeply explore signaling within and around the tumor, and also how it varied across the sample.

Molecular subtyping has traditionally been performed using RNA expression profiling of bulk tissues from MIBC samples(*8, 9, 16*). When compared to matched bulk profiling by RNA sequencing, we were able to robustly measure MIBC subtype markers with high predictive power in either the RNA or protein GeoMx assays. This demonstrates that from a single slide, it may be possible to capture and classify samples into subtype. Contamination of surrounding tissue may impact robustness of molecular classification. In our GeoMx experiment, this is avoided directly using auto-segmentation, allowing the investigation of compartment specific expression without having to manipulate the tissue or rely on pathological identification of tumor boundaries. Indeed, we noticed that stromal AOIs had significantly worse predictive performance, and application of auto-segmentation or strategic ROI placement is necessary for robust classification.

Furthermore, we observed little evidence of intra-tumor heterogeneity in molecular subtype markers. This is in contrast to recently published work suggesting that some tumors exhibit a high degree of intra-tumoral heterogeneity(*35, 36*). However, we note this difference may be due to definitions of subtype applied, the profiling strategy of this and other studies, and the small numbers of samples tested in each study. As our study was limited to 25 samples, we focused on detecting the differences between luminal and basal type tumors, as we would not be powered for further stratifications of the disease. We did note enrichment of TP53 mutations in specific luminal samples, however, without additional markers on the RNA-ISH panel we were unable to more deeply stratify samples.

In addition to subtype, the TME of MIBC is a critical component to patient outcome. We recapitulated the previously reported T-cell exclusion phenotype which was described in tumors expressing high levels of PPARγ(*6*). Specific exclusion of immune cell markers in the tumor compartment of PPARγ^high^ samples was observed across nearly all immune markers present on the panels in both the protein and ISH GeoMx studies. By directly measuring expression from stromal compartments, we were also able to assess whether the stromal immune composition of the PPARγ^high^ tumors were qualitatively different from PPARγ^low^ tumors. We found little evidence of differential expression between tumor based on PPARγ status, and genes that we did find to be differentially expressed were not generally immune-cell related. This supports the potential to improve ICI response by targeting the PPARγ pathway in these tumors, as this may remove the barrier to immune cell invasion in these tumors.

By spatially resolving the source of expression in a tissue, we were able to directly determine whether bulk profiling of tissues reflects a specific compartment within the sample, and identify novel interaction networks within and between tissue compartments. The bulk expression profiles were most similar to the tumor compartment, which may be due to the hyperproliferative nature of tumor cells. However, interrogation of other tissue types will be necessary to see if this results in a phenomenon that is broadly observed or specific to MIBC. *In silico* analysis of the TCGA suggest some tumor types may have more stromal signaling(*31*). Our findings suggest that even in indications with a high stromal content measured by bulk profiling, these studies may underestimate the role that the TME plays in tumor development by underrepresenting expression from the stroma itself. The observation in TCGA analysis that bladder cancers had lower tumor purity than many tumor types likely reflect the highly infiltrated phenotype of PPARγ^low^ tumors. In contrast, by directly measuring the stroma, we can begin to understand interactions between the compartments within that tissue. It is currently unknown if stromal signaling within MIBC may be related to tumor evolution and how to best target these tumors with therapeutics. The correlation analysis we employed suggests stromal expression of β-catenin may also be related to exclusion of immune cells in addition to tumor intrinsic PPARγ expression. While the functional role of β-catenin expression the stroma is unclear, these findings are reminiscent of work demonstrating the role of this pathway in immune cell exclusion in melanoma (*37*).

Beyond the key findings within MIBC, this study demonstrates the technical feasibility of using two sections of FFPE tissue per sample to obtain a highly nuanced view into the tumor-immune landscape. While this study was performed on archival tissue blocks, it provides a proof of principle that this technology can be applied to FFPE samples with limited material. Other technologies which address both location and multiplexing of analytes have distinct limitations in their ability to impact translational applications to clinically relevant samples such as degradation of signal during multiplexing, technical variability, and cumbersome downstream data analysis. Moreover, multiplexing of RNA probes in FFPE tissues has been a hurdle which few technologies have overcome. Technologies which offer the potential of higher multiplexing are limited to fresh-frozen tissues (25–27), and previously only low plex ISH approaches were compatible with the FFPE tissue which are most commonly available from the clinic. GeoMx RNA-ISH profiling offers a solution for high plex exploratory research on FFPE tissue that could be translated to low plex clinical assays on a diagnostic platform.

As GeoMx represents a complimentary approach for single-plex or multiplexed IHC and ISH, there are some differences between this platform and conventional profiling platforms that should be taken into consideration during experimental design. First, GeoMx profiling measures target abundance in an IF defined areas, which enables profiling of cell populations rather than individual cells. Second, as target quantification is mediated by direct probe hybridization, the platform is not capable of measuring somatic mutation, and assays capturing oncogenic mutations and TMB remain a complementary approach. This was demonstrated in our study, as one PPARγ^high^ sample, 10182, had higher TMB and a specific SMAD4 mutation (T338I) that may explain the immune infiltration observed in this tumor, given the known impact of TGFβ signaling in immune infiltration and ICI response(*34*). Finally, GeoMx profiling does not require cycling to achieve high plex, where stripping and re-probing with new antibodies tends to degrade signal over time. Rather the quantification of all targets is performed simultaneously and maintains their relative abundance.

While this study highlights several key advantages of performing GeoMx on clinically relevant samples, we note that there are several limitations to our study. As we only tested 157 genes in the RNA-ISH and 40 antibodies, there are a significant number of pathways or targets of interest that were not measured. Also, while this manuscript presents some preliminary strategies to capture the cell-type diversity and interactions between tumor and stromal compartments, statistical methodologies will need to be adapted for the unique characteristics of the data being generated on this platform. For example, methodologies aimed at cell-type deconvolution, including CIBERSORT (*38, 39*), TIMeR (*40*), and MuSIC (*41*), have been shown to capture the cellular composition of bulk-profiled samples. These algorithms may need to be adapted for spatial profiling by incorporating AOI-specific information such as AOI size or number of cells. Other spatially resolved platforms have started to propose modification to differential expression analysis to leverage the spatial data captured(*42*). Due to the sparse collection strategy employed such methods may need to include additional caveats to be applied to GeoMx profiling data.

With rapid advances in the ability to profile tissues in a spatially resolved manner, we have found that GeoMx provides a robust, targeted platform for profiling tissues that is readily geared towards exploratory studies. Incorporating it with multiple platforms in this study allowed us to extend beyond previous foundational studies, and deeply profile the tumor-stroma interaction in MIBC. Such profiling strategies have a clear benefit for translational researchers allowing spatial profiling while maintaining high-plex quantification of FFPE samples.

## Materials and Methods

### Tissue Procurement

25 formalin-fixed, paraffin-embedded (FFPE) samples from bladder cancer patients were purchased from Proteogenex and banked at Eisai (Andover, MA). Informed consent was obtained from all patients by the Russian Oncological Research Center n.a. N.N. Blokhin Rams Ethics Committee.

### Bulk RNA profiling by IO 360 and RNA sequencing

For each sample, 5 of slides were processed using the Qiagen RNeasy FFPE Kit to extract total RNA from 10 µM FFPE tissue sections. RNA was profiled using the NanoString nCounter IO 360 panel, using 100 ng of RNA per sample profiled using the nCounter Max Profiler. Expression of all probes was normalized based on the housekeeping probes included in the panel such that the geometric mean of expression of the housekeeping genes for a sample was centered at a constant value of 7 in log_2_ expression space for all samples. 100 ng RNA was also profiled using TruSeq RNA access Library Prep Kit on an Illumina Nextseq to a depth of 200 million reads with a minimum of 10 million reads. RNA-sequencing reads were mapped using the STAR aligner (*43*) and quantified using Kallisto (*44*).

### Immunohistochemistry and In Situ hybridization

Immunohistochemistry Primary antibodies include granzyme B clone 11F1 (Leica Biosystems), PPARγ clone C26H12 (Cell Signaling Technology), CD8 clone SP57 (Ventana Medical Systems) and PD-L1 clone 22C3 (Agilent Technologies). The CD8 assay was performed on the Ventana automated immunostainer BenchMark ULTRA. The PDL1 assay was performed using EnVision FLEX HRP visualization system on DAKO Autostainer Link 48 platform. PDL1 staining was scored by a board-certified MD-pathologist at Cancer Genetics Inc. PPARg RNA-ISH was performed using the RNAscope technology in the service lab at Advanced Cell Diagnostics. CD8 IHC, granzyme B IHC and PPARG RNA-ISH staining was quantified using the digital pathology software Halo (Indica Labs). Scanned images, pathologist scores and interpretations were available to H3 Biomedicine Inc., for review and data analysis.

### Mutation profiling and TMB detection

DNA was extracted from a single FFPE slide per sample and 40 ng of DNA was used as input for sequencing on the TruSight Oncology 500 (TSO-500) Illumina targeted mutation profiling panel. Estimation of tumor mutational burden (TMB) was performed based on the number of estimated somatic mutations for either non-synonymous variants or all somatic alterations detected. Potential oncogenic impact of specific mutations was inferred using Cscape (*33*).

### GeoMx profiling of Antibody & RNA-ISH expression

Slides were prepared by incubating with a cocktail of either antibodies or ISH probes conjugated to photocleavage reporter tags. After incubation with reporter reagents, slides were incubated with IF-conjugated Pan-CK and CD3 probes, as well as a DNA stain. Samples were imaged on the GeoMx digital spatial profiler platform, wherein an IF microscope was used to guide live image selection of regions of interest placement. AOIs were created from ROI images using PanCK IF staining, creating PanCK+ and PanCK-segments. UV light was shown onto the tissue based on the segments defined by PanCK staining using a digital micromirror array, and tag oligos were collected for downstream quantification. Antibody conjugated tags were profiled using the nCounter Flex system, while the RNA-ISH probes were quantified using an Illumina NextSeq. Sample preparations, AOI mask creation, and quantification is described in detail in the supplementary Methods.

### Bioinformatics and Statistical Analysis

For RNA-sequencing data, expression analysis was performed using Limma-voom(*45*) and pathway analysis was performed using Gene Set Enrichment Analysis(*32*). Reference gene sets were obtained from the Molecular Signatures Databases (MSigDB) (*32*), and fisher’s exact test was used to test for the significance of the difference between the enriched genes and all the genes in the ISH panel for each of the signatures evaluated.

We defined the molecular subtype of “basal” vs “luminal” from the RNA-sequencing data using a previously published signature(*9*). Samples were classified into PPARγ^low^ and PPARγ^high^ groups based on the median mRNA expression of PPARγ based on NanoString counts, after normalization. After normalization of the IO360 expression data, previously described signatures(*21, 22, 46*) were calculated across all samples. Briefly, expression signatures were calculated for each sample the using a linear model such that:

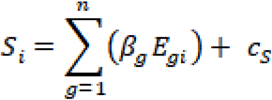

Where the signature score, *S*, for sample, *i*, is the sum of the expression of each gene for a sample, *E*_*gi*_, multiplied by predefined weights for each signature gene, *β*_*g*_, and a constant for each signature, *c*_*S*_. Scores were calculated and reported in log_2_ transformed expression space.

Correlation between analytes were calculated using either Spearman correlation coefficient (ρ) or the Pearson correlation coefficient (R^2^) as appropriate. Unsupervised hierarchical clustering was performed using complete linkage and Euclidean distance for clustering Z-scores of expressions of a given probe or analyte.

Differential protein expression from GeoMx data between phenotypes of interest was calculated using a mixed effect model allowing for random intercepts and slopes described as:

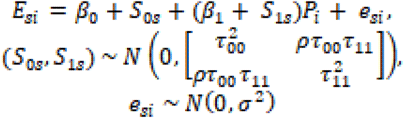

Where the protein expression of a probe, *E*_*si*_, is tested against a phenotype, *P*_*i*_ (e.g. AOI compartment), allowing for random intercepts and slopes modeling patient sample, *S*, to account for multiple AOIs tested within a patient slide. P-values for the mixed effect models were calculated using Satterthwaite’s approximation for degrees of freedom(*47–49*).

P-values from statistical models were adjusted for multiple testing using the Benjamini-Hochberg false-discovery rate (FDR)(*50*).

## Supplementary Materials

### Materials and Methods

Table S1. Clinical and molecular characteristics of MIBC cohort samples.

Table S2. RNA-ISH AOI characteristics and QC metrics.

Table S3. Spatial differential expression and compartment interaction analysis

Data File S1. GeoMx Antibody data used in analysis

Data File S2. GeoMx RNA-ISH data used in analysis

Data File S3. Bulk expression signature analysis

Fig. S1. ROI placement on guide IF images

Fig. S2. QC of GeoMx profiling

Fig. S3. Relationship between bulk and GeoMx expression

Fig. S4. IHC quantification of CD8 and GZMB

Fig. S5. AOI expression of CD3 antibody

## Acknowledgments

We thank Alessandra Cusano, Lihua Yu, and Ping Zhu for helping review the manuscript.

## Author contributions

JWR, ZZ performed data analysis and prepared the manuscript; DMZ, JG coordinated and ran GeoMx profiling; ZKN created and ran the RNA-ISH sequencing pipeline; SD, MK, PK, VR procured FFPE samples and coordinated mutation profiling, RNA sequencing, and IHC profiling; SW, MH, JB, PK, VR conceived of study design and managed study execution

## Data and materials availability

Processed data files are provided as supplemental material. Original nCounter expression files have been uploaded to GEO, and RNA sequencing files have been uploaded to the SRA. High resolution image files may be made available through an MTA.

## Supplementary Methods

### Sample preparation and readout of protein Digital Spatial Profiling

For multiplex antibody analysis, a cocktail of 40 primary antibodies (Supplementary Data File 1), each with a unique, UV photocleavable indexing oligo, and 2 fluorescent markers (Pan-CK and CD3) were added to a 5 µm FFPE slide at 4°C overnight. Pan-CK and CD3 were used to identify tumor cells and T cells respectively. Syto83 (nuclear DNA stain) was added to the FFPE tissues for 5 minutes at room temperature. After washing off the extra antibodies in TBST, the tissue slides were placed on the stage of an inverted microscope. A custom gasket was then clamped onto each slide, allowing the tissue to be submerged in 1.5 mL of buffer solution. Under the microscope, wide field fluorescence imaging was performed with epi-illumination from visible LED light engine. 20x images were stitched together to yield a high-resolution image of the tissue area of interest. Two of the 25 samples (10171 and 10230) were omitted from further processing due to poor tissue attachment. Of the remaining 23 samples, regions of interest (ROIs) were then selected based on the fluorescence information and sequentially processed by the microscope automation. Tumor and stromal segmentations were processed for each ROI based on Pan-CK+ and - using image J scripts as described below. UV LED light was collimated to be reflected from the digital micromirror device (DMD) surface into the microscope objective and focused at the sample tissue with paired segmentation mask (described below). A microcapillary tip connected to a syringe pump primed with buffer solution was moved to each ROI and collected oligos released from the tissue by UV irradiation. These oligos were further quantitated using Nanostring GeoMx™ Digital Spatial Profiler.

Hybridization of cleaved indexing oligos to NanoString barcoded sequences was performed using the nCounter PlexSet reagents. Oligos were denatured at 95°C for 3 to 5 minutes and placed on ice for 2 minutes. A master mix was created by adding 70 μL of hybridization buffer and in situ capture probes (ICP) to the PlexSet tube (A-G). A 7μL aliquot of master mix was added to each well in 96-well plate. And denatured protein samples were added to the well and the volume of each well was brought to a final volume of 15 μL with DEPC treated water. Hybridizations were performed at 65°C overnight in a thermocycler. After hybridization, samples were pooled by column and processed using the nCounter Prep Station and Digital Analyzer as per manufacturer instructions.

Digital counts from tags corresponding to protein probes were analyzed as follows: raw counts were first normalized with internal spike-in controls to account for system variation. These normalized values were then normalized by adjusting for the area of sample illuminated by UV and log_2_ transformed prior to analysis. Antibodies with mean expression below the mean of the highest IgG control antibody were excluded from analysis.

### Sample preparation for ISH Digital Spatial Profiling

For in situ hybridization, 5 µm FFPE sections were freshly cut and mounted onto positive charged slides were baked, deparaffinized, rehydrated in ethanol and washed in PBS using Leica Bond Rxm system. Targets were retrieved for 20 minutes in Leica Epitope Retrieval Solution 2, EDTA PH 9.0 buffer at 100°C. Tissues were washed with PBS and incubated with 1ug/ml proteinase K in PBS for 15 mins at 37°C and washed with PBS. After tissues were removed from the Leica Bond, they were covered with Hybrislip covers (Grace BioLabs) and incubated overnight at 37°C with hybridization solutions containing 4 nM of each RNA detection probe; 0.1 mg/mL salmon sperm DNA (Sigma-Aldrich, D7656); 2.5% dextran sulfate (Sigma-Aldrich, 67578-5G); 0.2% BSA (ThermoFisher, 37525); 40% deionized formamide (Ambion, AM9344); 2X SSC (Sigma, S6639). After overnight incubation, HybriSlip covers were gently removed by soaking in 2X SSC + 0.1% Tween 20 and then washed for 5 minutes in 2X SSC. Two 25-minute stringent washes were performed in 50% formamide in 2X SSC at 37°C. Tissues were washed twice for 2 minutes each in 2X SSC, and then were blocked with a Nanostring blocking buffer for 30 minutes at room temperature in a humidity chamber. SYTO 13 and fluorescently-conjugated antibodies targeting Pan-CK and CD3 in blocking buffer were applied for 1 hour at room temperature, then washed three times for 5 minutes each in fresh 2X SSC. Three of the 25 samples (10167, 10171 and 10230) were omitted from further processing due to poor tissue attachment. Oligos were cleaved and collected using Nanostring GeoMx™ Digital Spatial Microscope as described above.

### Custom UV illumination mask creation

For each sample, 6 AOIs were selected and the oligonucleotide tags from stroma vs tumor areas were collected and analyzed separately by creating a Pan-CK+ and Pan-CK-UV illumination mask for each AOI. Custom masks were created in ImageJ to define custom regions of interest for UV illumination. These custom masks were used by the DMD device to determine which mirrors would be utilized to direct UV light to tumor and stromal segmentations. For each region of interest, 20x stacked tiff images with four fluorescent channels was split into four separate fluorescent images for custom thresholding. To make tumor segmentation, the image with Pan-CK channel was thresholded manually to match the Pan-CK staining pattern and converted to a binary mask. Remove fluorescent noise by copying a mask generated using “Analyze Particles” with settings size = 0-700 pixels, circularity = 0.35-1.00, and show = masks, to Pan-CK binary mask. “Fill holes” was proceeded to fill the nucleus of the tumor cells. “Invert LUT” is to make sure Pan-CK positive region could be UV illuminated. This image was saved as tumor segmentation. Stromal segmentation was generated by inverting tumor segmentation following by being dilated 3 times. Segmentation size is defaulted with 666×666 µm box. However, in some ROIs, only center region is designed to be UV radiation exposed. In this case, a centered 665 µm diameter circle mask could be added to tumor or stromal segmentations by using “Clear outside” function.

### RNA ISH profiling readout by NGS

PCR was performed on photoreleased oligos collected from each well (corresponding to one AOI) to add Illumina adapter sequences and unique i5 and i7 sample indices. Each PCR reaction used 4 uL of DSP photocleaved oligos, 1 uL of unique-dual indexing PCR primers, 2 uL of Nanostring 5X PCR Master Mix, and 3 uL PCR-grade water. Thermocycling conditions were 37C for 30 min, 50C for 10 min, 95C for 3 min; 18 cycles of 95C for 15sec, 65C for 1min, 68C for 30 sec; and 68C 5 min. PCR reactions were pooled and purified twice using AMPure XP beads (Beckman Coulter) according to manufacturer protocol. Pooled libraries were sequenced at 2×38 base pairs and with the dual index workflow on an Illumina NextSeq to generate 306M raw reads.

For data from the NextSeq instrument, FASTQ files from multiple lanes were merged to generate single files for processing and insure proper removal of PCR duplicates later in the pipeline. Illumina adapter sequences were trimmed using Trim Galore (version 0.4.5) with a minimum base pair overlap stringency of four bases and a base quality threshold of 20. Paired end reads were stitched using Paired-End reAd mergeR (PEAR, version 0.9.10) specifying a minimum stitched read length of 24bp and a maximum stitched read length of 28bp. The 14bp UMI sequence was extracted from the stitched FASTQ files from the 5’ end of the sequence reads using umi tools (version 0.5.3). The fastq files with extracted UMIs were then aligned to a genome containing the 12bp reference sequence tags using bowtie2 (version 2.3.4.1) in end-to-end mode with a seed length of four. Using a custom python function, the generated SAM files were split in to multiple SAM files based on the tag to which they aligned to limit memory usage when removing PCR duplicates. The split SAM files were converted to bam files, sorted, and index using samtools (version 1.9) with the import, sort, and index options respectively. PCR duplicates were removed from the sorted and indexed bam files using the dedup command from umi tools with an edit distance threshold of three. An edit distance threshold of three was used because using a threshold of one quarter the length of the UMI has been demonstrated to be a conservative threshold. Using custom python functions, the SAM files with PCR duplicates removed were merged for each sample and used to generate digital counts of the tags.

Because each target transcript was counted using multiple probes and tags, outlier counts were removed before generating a consensus count for each target. Outlier tags were identified as those with counts 90% below the mean of the probe group in at least 20% of the AOIs analyzed and completely removed them from the analysis. Subsequently we removed tags from the analysis if they were flagged as outliers in at least 20% of the AOIs analyzed. This was done using the Rosner Test if there were at least 10 probes for the target (k = 0.2 * Number of Probes, alpha = 0.01), or the Grubbs test if there were less than 10 probes for the target. Probes flagged as outliers in less than 20% of the AOIs analyzed were only removed from the analysis for the AOIs in which they were flagged. Count reported for each target transcript were calculated as the mean of the remaining probes.

The counts for each target transcript were then normalized to the count of the house keeper genes (C1orf43, GPI, OAZ1, POLR2A, PSMB2, RAB7A, SDHA, SNRPD3, TBC1D10B, TPM4, TUBB, UBB). The geometric mean of the house keeper gene counts was calculated for each AOI. These geometric means were then divided by the geometric mean of the geometric mean of the house keepers to generate a normalization factor for each AOI. The counts of the transcripts in each AOI were than multiplied by the associated normalization factor. Expression values were log_2_ transformed prior to analysis.

---

**Supplementary Figure 1.**
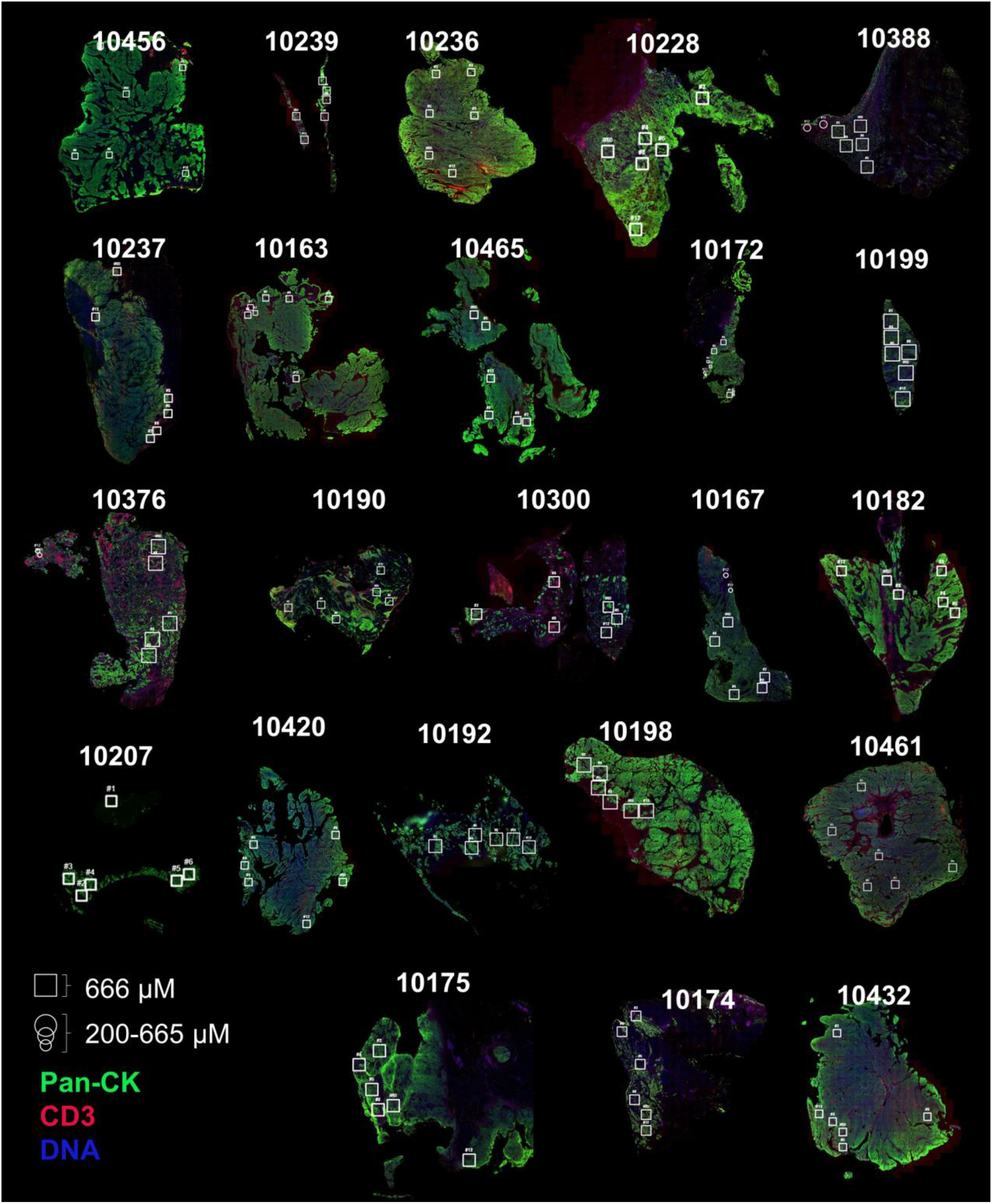
ROI placement on guide IF images. Whole-slide IF images are shown for each sample indicating the placement of ROIs throughout the tissue. Each slide is scaled independently, though square ROIs are consistently sized, at 666 uM per ROI. Circular ROIs present in some samples were not used during analysis, as they varied in size and cell type composition.

**Supplementary Figure 2.**
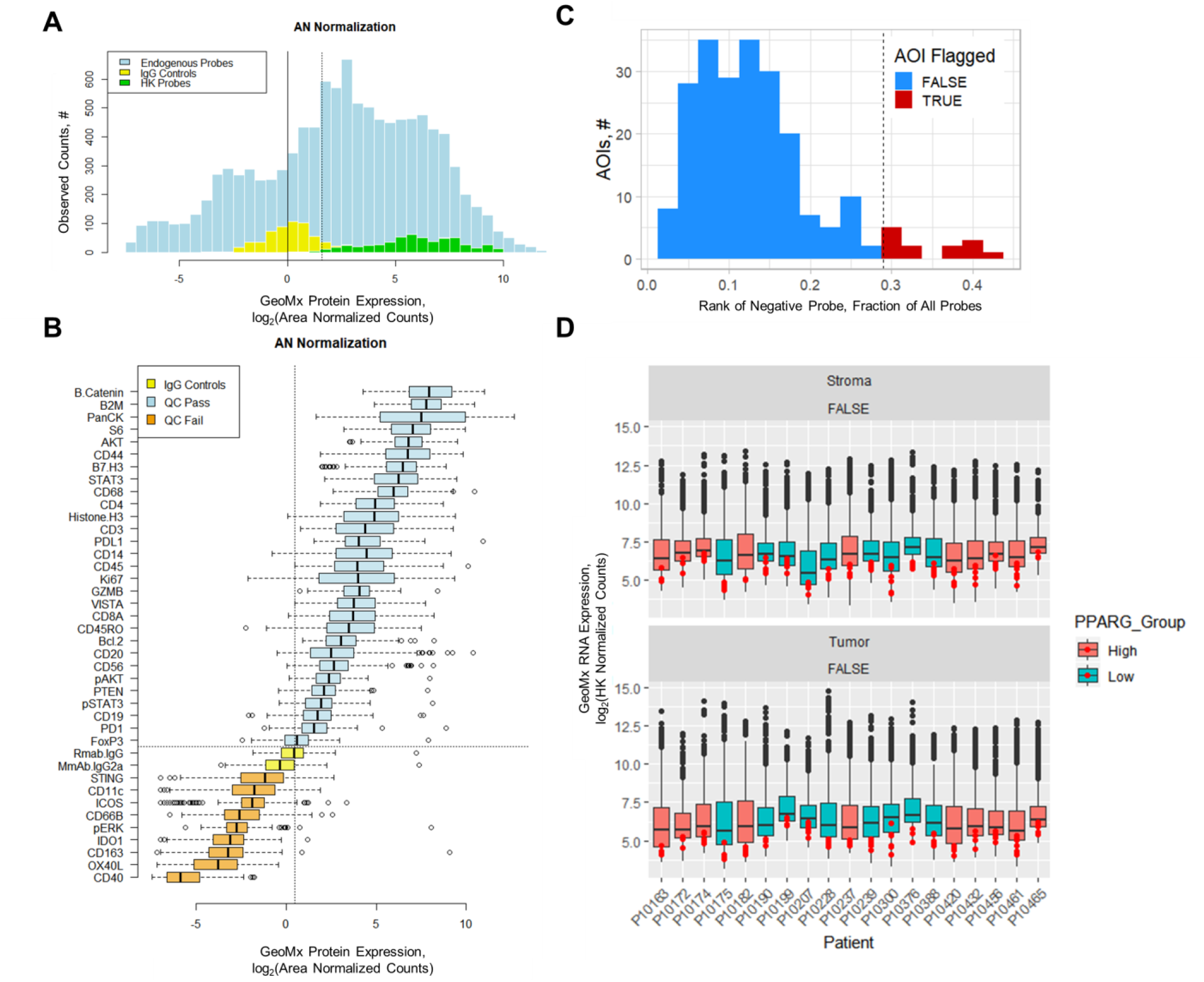
QC of GeoMx profiling. **(A)** Distribution of counts produced across probes from different classes of antibodies. HK probes include Histone H3, and S6. Control probes are Rabbit IgG1 (Rmab.IgG) and Mouse IgG2A (MmAb.IgG2a). All other probes are for endogenous expression detection. **(B)** Individual probe distribution and QC. Probes falling below the median expression of the Rabbit IgG1 probe were excluded from analysis due to poor signal. **(C)** Distribution of the rank of the negative control probes from RNA-ISH profiling across AOIs output from RNA-sequencing pipeline. AOIs with high ranking negative probes (e.g. high background samples. > Mean + 2σ), were removed from analysis (n = 13). **(D)** Distribution of RNA-ISH expression across all AOIs for all genes measured, aggregated by patient. Red dots indicate the negative control probe expression for each AOI within a patient block.

**Supplementary Figure 3.**
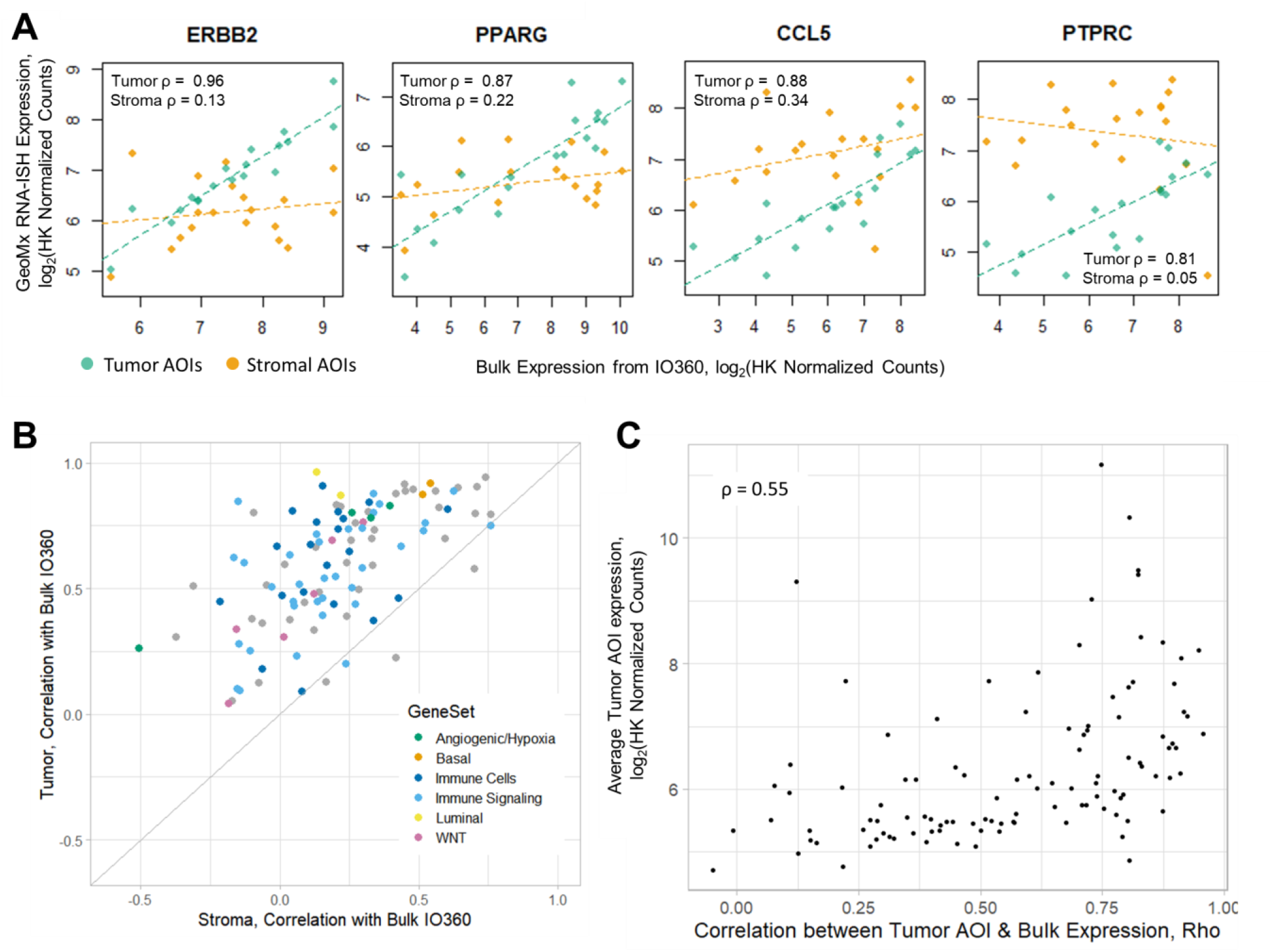
Relationship between bulk and GeoMx expression. **(A)** Example expression patterns for tumor and immune cell related genes. Correlation between average tumor AOI or average stromal AOI expression shown compared to bulk profiling by IO360. **(B)** Correlation of individual genes with either stroma (x-axis) or tumor (y-axis) expression. Probe colored by gene set membership **(C)** Expression of tumor AOIs is weakly correlated with how well correlated a given probe is measured in tumor AOIs compared to bulk expression.

**Supplementary Figure 4.**
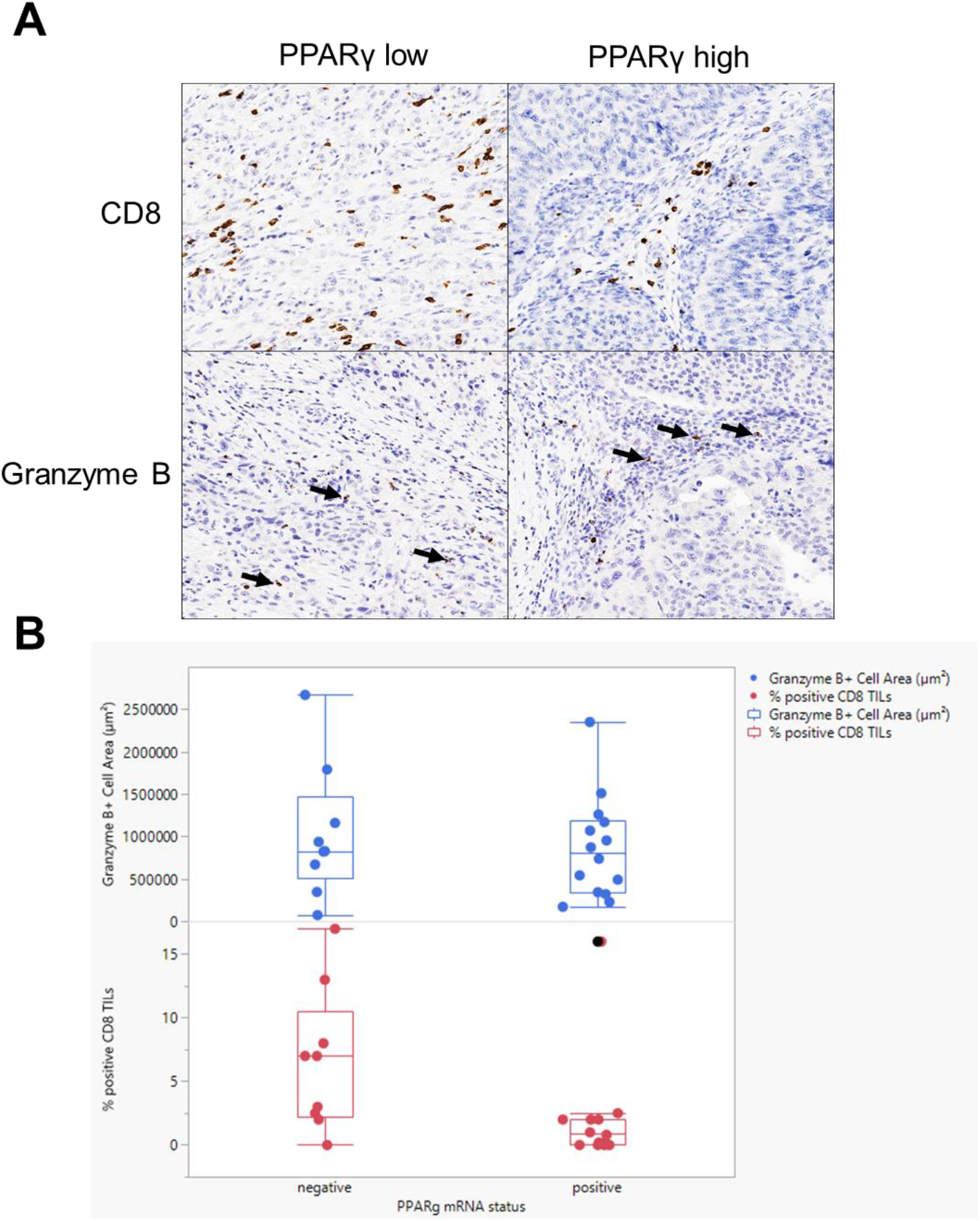
IHC Quantification of CD8 and GZMB. **(A)** Example staining of CD8 and Granzyme B by IHC in a representative PPARg negative and positive tumor. Arrows indicate GZMB+ positive cells. **(B)** Quantification of samples stained with CD8 and GZMB across the cohort of samples based on PPARG status.

**Supplementary Figure 5.**
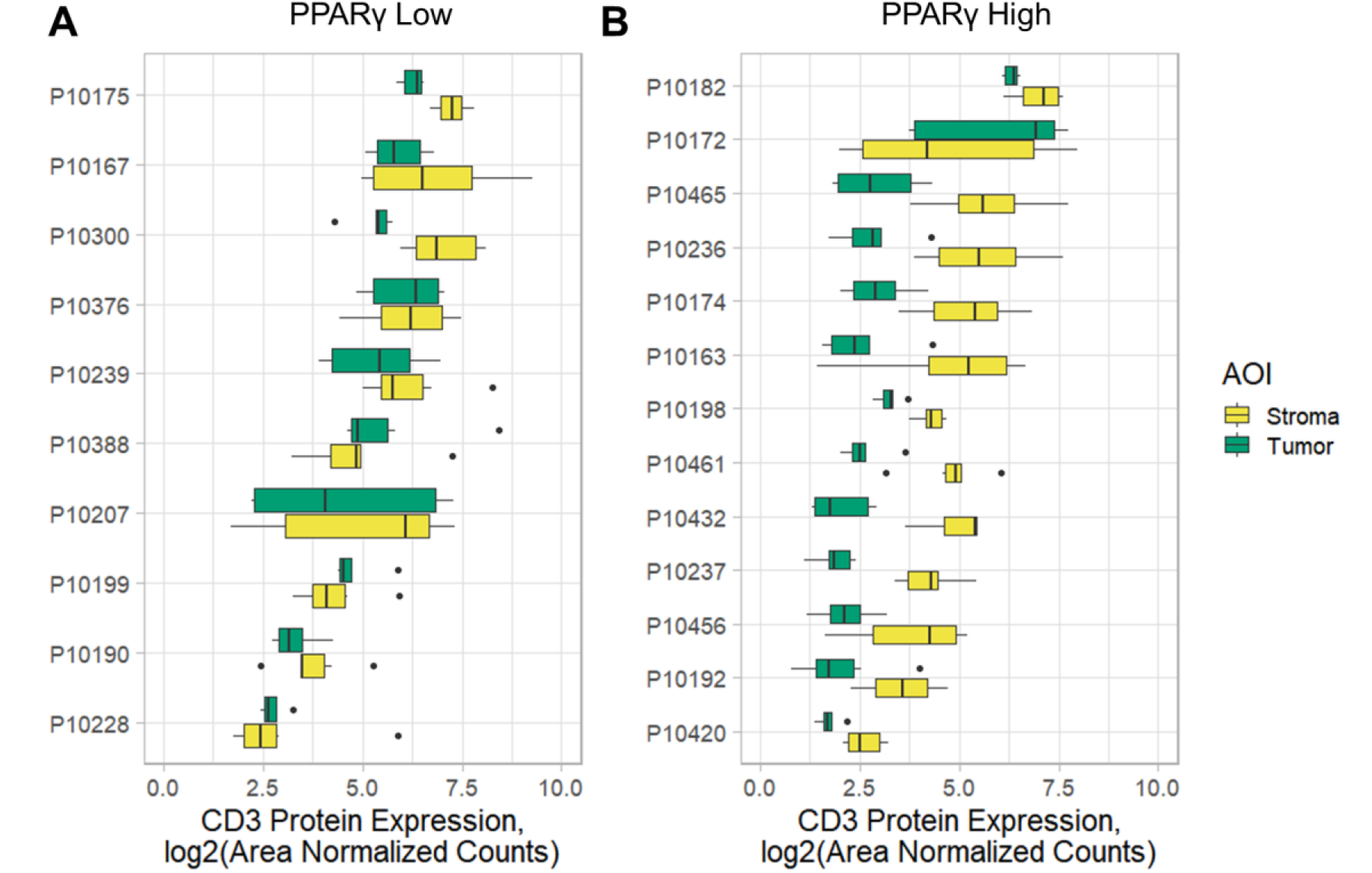
AOI expression of CD3 antibody. **(A-B)** Patient level expression of CD3 for either **(A)** PPARγ^low^ or **(B)** PPARγ^high^ samples.

